# Phase separation of TPX2 enhances and spatially coordinates microtubule nucleation

**DOI:** 10.1101/668426

**Authors:** Matthew R. King, Sabine Petry

## Abstract

Phase separation of substrates and effectors is proposed to enhance biological reaction rates and efficiency. TPX2 is an effector of microtubule nucleation in spindles, and functions with the substrate tubulin by an unknown mechanism. Here, we show that TPX2 phase separates into a co-condensate with tubulin, which mediates microtubule nucleation *in vitro* and in isolated cytosol. TPX2-tubulin co-condensation preferentially occurs on pre-existing microtubules at the endogenous and physiologically relevant concentration of TPX2. Truncation and chimera versions of TPX2 directly demonstrate that TPX2-tubulin co-condensation enhances the efficiency of TPX2-mediated microtubule nucleation. Finally, the known inhibitor of TPX2, the importin-α/β heterodimer, regulates both co-condensation and activity. Our study demonstrates how regulated phase separation can simultaneously enhance reaction efficiency and spatially coordinate microtubule nucleation, which may facilitate rapid and accurate spindle formation.

## Introduction

The microtubule (MT) cytoskeleton organizes the interior of the cell, determines cell shape, and segregates chromosomes. Underlying its timely and accurate formation are multiple MT nucleation pathways from various cellular locations that need to be turned on at the right cell cycle stage. Only few pathway-specific MT nucleation effectors are known and their molecular mechanisms remain poorly understood (Petry, 2016; Tovey and Conduit, 2018). At the same time, pioneering *in vitro* studies have implicated a role for liquid-liquid phase separation (LLPS) of proteins in cytoskeletal assembly (Hernández-Vega et al., 2017; Jiang et al., 2015; Li et al., 2012; Su et al., 2016; Woodruff et al., 2017), but the exact physiological contribution remains unclear. More generally, many proteins have been shown to undergo LLPS *in vitro*, while functional roles of LLPS in cells remain to be discovered (Banani et al., 2017; Shin and Brangwynne, 2017).

Branching MT nucleation is a recently identified pathway during which new MTs nucleate along the lattice of pre-existing ones (Petry et al., 2013). It exponentially increases MT numbers while preserving their polarity and is critical for rapid and accurate spindle assembly (Decker et al., 2018; Kaye et al., 2018; Petry et al., 2013). Branching MT nucleation requires the universal MT nucleator module, consisting of the γ-Tubulin Ring Complex (γ-TuRC) (Oegema et al., 1999; Zheng et al., 1995) and its recently discovered co-factor XMAP215 (Flor-Parra et al., 2018; Gunzelmann et al., 2018; Thawani et al., 2018), as well as the protein complex augmin that directly recruits γ-TuRC along the length of a pre-existing MT (Song et al., 2018). Branching MT nucleation is initiated by the microtubule associated protein TPX2 (Petry et al., 2013), which has been proposed to activate γ-TuRC via TPX2’s C-terminal domain (Alfaro-Aco et al., 2017; Scrofani et al., 2015). *In vitro*, TPX2 can directly generate MTs from tubulin via its N-terminal domain (Roostalu et al., 2015; Schatz et al., 2003), but this domain is dispensable for MT nucleation in cytosol (Alfaro-Aco et al., 2017; Brunet et al., 2004). Understanding how TPX2 stimulates MT nucleation could help pioneer how a specific MT nucleation pathway is turned on in order to build cellular MT structures such as the mitotic spindle.

Here, we show that TPX2 undergoes phase separation to form a co-condensate with tubulin at its endogenous and physiologically relevant concentration in *Xenopus* egg cytosol. The co-condensation of TPX2 and tubulin occurs on microtubules and thus helps to specifically promote MT nucleation from pre-existing MTs and enhance MT nucleation rates in cytosol. Lastly, importins regulate this process by inhibiting the formation of co-condensates. Collectively, these data provide a molecular mechanism for TPX2 function which is not only critical to explain spindle assembly, but also demonstrates that phase separation can spatially coordinate reactions and enhance reaction kinetics in a physiological context.

## Results

### TPX2 forms a co-condensate with tubulin in vitro and in cytosol

When characterizing TPX2, we noticed features of known phase separating proteins: a disordered N-terminus and a more ordered C-terminus with potentially multivalent alpha-helical regions (Alfaro-Aco et al., 2017) (Fig. 1A). When either GFP-tagged or untagged TPX2 in high salt buffer was introduced to physiological salt levels, spherical condensates formed (Fig. 1B). These condensates fulfill several criteria of LLPS: they fuse, exhibit salt- and concentration-dependent condensation, and show fluorescence recovery that saturates over time (Fig. S1A-C, Movie S1).

**Figure 1:**
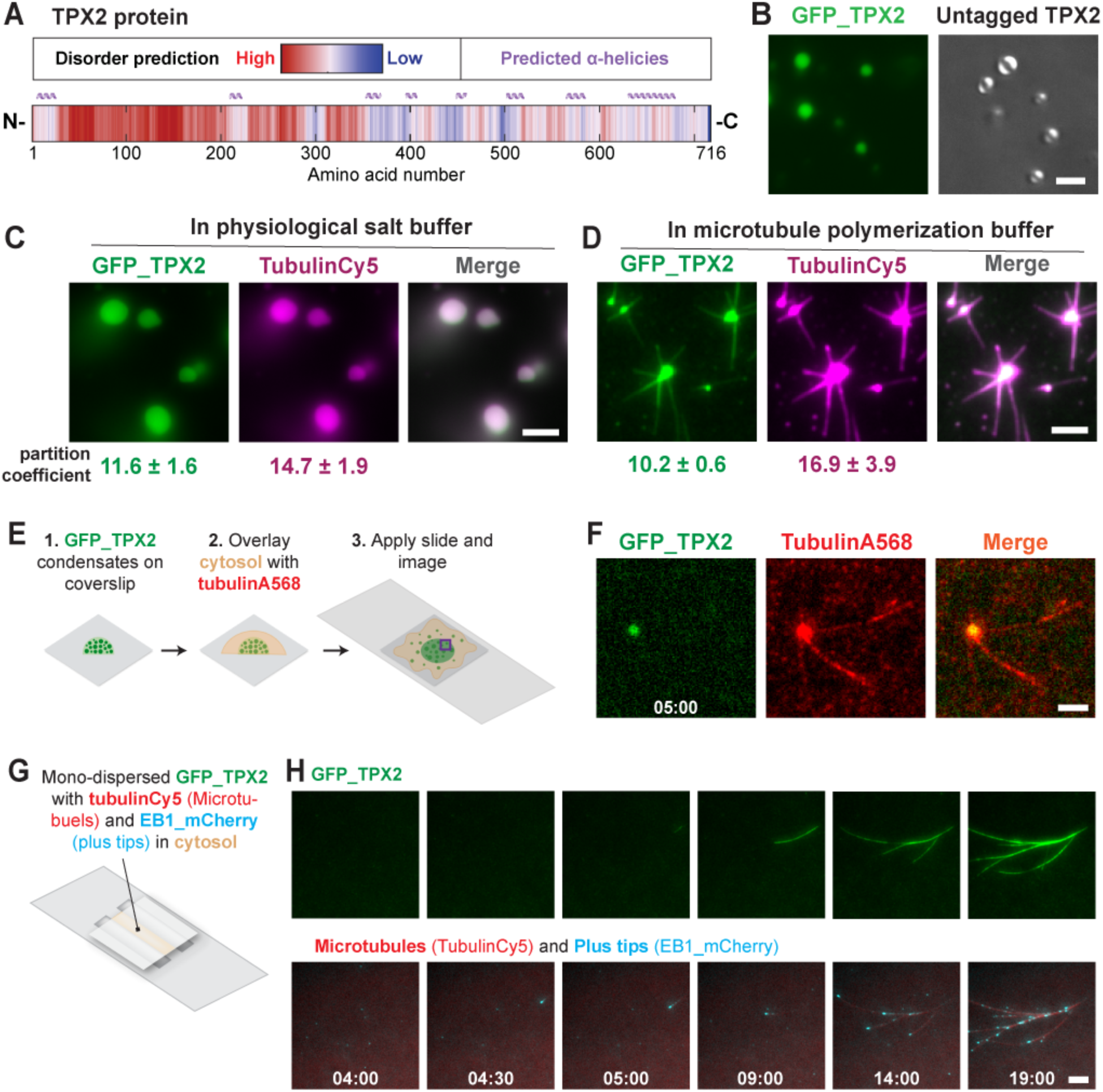
TPX2 forms a co-condensate with tubulin *in vitro* and in cytosol. **(A)** Intrinsic disorder and secondary structure predictions in TPX2. **(B)** Epifluorescent image of GFP-TPX2 condensates (See Movie S1) (left) and DIC image of untagged TPX2 condensates (right) **(C)** Epifluorescent image of GFP-TPX2 condensates prepared with Cy5-labeled tubulin (both at 4 µM) in a flow chamber (see figure S1D for control) **(D)** TIRF image of TPX2-Tubulin co-condensates (1 and 10 µM, respectively) prepared in MT polymerization buffer in a flow chamber, 18 minutes after reaction started. **(E)** Experimental set up for (F). Purple square roughly indicates area imaged. **(F)** Oblique-TIRF microscopy of GFP-TPX2 and tubulin (Alexa568-labeled) taken 5 minutes after reaction started (minutes:seconds). See also Figure S2 for additional time-points and additional treatments. **(G)** Experimental set up for (H) and (Fig. 2C): mono-dispersed GFP-TPX2 (500nM) and Alexa568-labeled tubulin mixed with *Xenopus* meiotic egg cytosol and imaged via oblique-TIRF microscopy in a flow chamber **(H)** GFP-TPX2 localization to growing MT-network imaged over time (minutes:seconds) in cytosol (see Movie S2). Microtubules are labeled red and growing plus-tips are blue. All scale bars are 3 µm and TPX2 was [2µM] unless indicated. Partition coefficient values are mean with ±1 standard deviation (SD). Data shown are representative of at least three experimental replicates.

We hypothesized that TPX2 may interact with tubulin dimers as a co-condensate because the two do not interact as mono-dispersed proteins yet form ‘clusters’ that nucleate MTs both *in vitro* (Alfaro-Aco et al., 2017; Roostalu et al., 2015; Schatz et al., 2003) and in cells (Brunet et al., 2008; Ma et al., 2010; Tulu et al., 2006). Indeed, TPX2 forms a co-condensate with tubulin that generates MTs (Fig. 1C-D), forming an aster similar to those previously observed *in vitro* (Schatz et al., 2003). Importantly, TPX2 selectively co-condenses with tubulin but not with a protein of similar size and charge (Figure S1D), demonstrating that TPX2 and tubulin specifically from MT-nucleation competent co-condensates.

To investigate the function of TPX2-tubulin co-condensation in a physiological context, pre-formed TPX2 condensates were overlaid with meiotic *Xenopus laevis* egg cytosol containing soluble tubulin (Fig. 1E). TPX2 condensates selectively enriched tubulin from the isolated cytosol and generated branched MT networks (Fig. 1F, S2A). The tubulin signal in the condensates diminished as they generated branched MT networks (Fig. S2B), but not as a result of photobleaching (Fig. S2C). The physiological behavior of TPX2 to generate branched MT networks could only be observed with non-aged, liquid-like TPX2 condensates, but not with TPX2 condensates that had hardened after aging (Fig. S1C and S2D-F). The latter condensates still enriched tubulin and generated either aster-like MT arrays (when aged 15 min, Fig. S2E) or no visible MTs (when aged 30 min, Fig. S2F). In addition, we added mono-dispersed GFP-TPX2 to meiotic *Xenopus laevis* egg cytosol, which, in this reaction, stimulates the formation of branching MT nucleation and binds to emerging MTs (Fig. 1G-H, Movie S2). Collectively our observations demonstrate that TPX2 and tubulin can undergo LLPS to form a co-condensate capable of generating MTs *in vitro* and in cytosol. These data reconcile previous observations of TPX2-tubulin ‘clusters’ and provide a mechanistic framework for how the two proteins may functionally interact.

### TPX2-tubulin co-condensates form on MTs at the endogenous and physiologically relevant concentration of TPX2

To further investigate the functional significance of TPX2-tubulin co-condensation, we mapped the phase boundary of TPX2 via partition coefficient (Fig. S3A-B) and soluble pool measurements (Fig. S3C). Interestingly, tubulin lowers the concentration at which TPX2 phase separates from ∼200nM to ∼50nM, which, compellingly, corresponds to the estimated concentration range for endogenous TPX2 in *Xenopus laevis* egg cytosol of 30-100nM (Fig. S3A-C) (Wühr et al., 2014). Surprisingly, TPX2 localizes to microtubules at the much lower concentration of 1nM (Fig. 2A-B), which is 50-fold lower than the TPX2 phase boundary in the presence of tubulin in solution (Fig. S2A-E). This suggests that TPX2 prefers to bind to MTs over associating with itself or tubulin in solution. Interestingly, MT-localized TPX2 can still specifically recruit soluble tubulin along the length of MTs (Fig. 2A, S3F). Moreover, this occurs at the same concentration of 50nM TPX2 that corresponds to the phase boundary of TPX2-tubulin co-condensation in solution (Fig. 2B, S3D-E), indicating that TPX2-tubulin co-condensates form on MTs. Based on these results, we hypothesized that the only condition, at which TPX2-tubulin co-condensates can form in solution and not on MTs is if MT formation is prevented. Indeed, when MT polymerization in cytosol was inhibited via nocodazole, mono-dispersed TPX2 and tubulin formed small, spherical, and highly mobile TPX2-tubulin co-condensates (Fig. 2C, Movie S3). These are reminiscent of nocodazole-induced TPX2 ‘puncta’ previously observed in cells (Ma et al., 2010; Tulu et al., 2006). In sum, our evidence suggests that TPX2-tubulin co-condensates form on MTs, rather than in solution.

**Figure 2:**
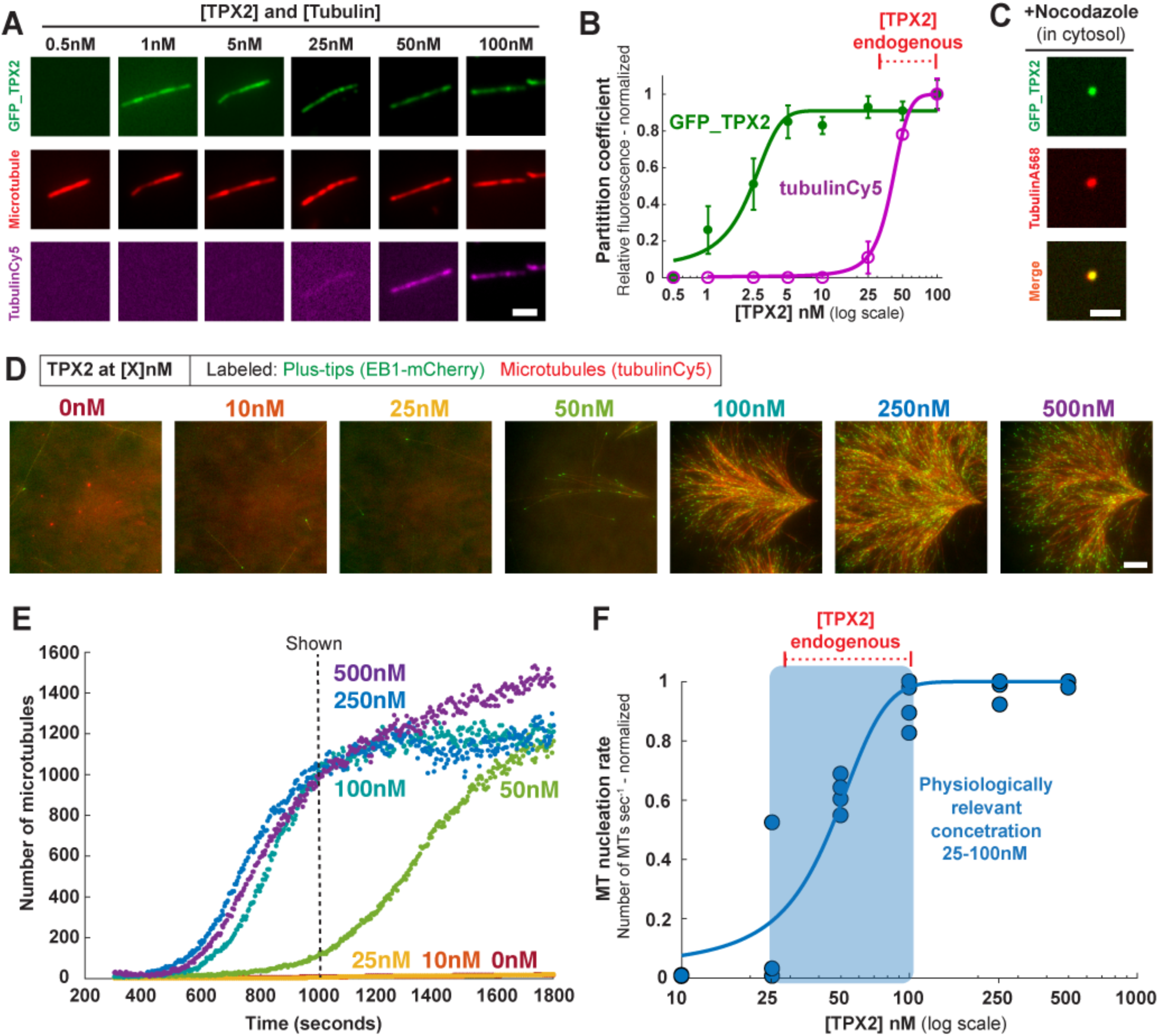
TPX2-tubulin co-condensates form on MTs at the endogenous and physiologically relevant concentration of TPX2. **(A)** TIRF images (contrast-optimized) and **(B)** quantification of GFP-TPX2 and Cy5-labeled tubulin (at indicated concentrations, both proteins equimolar) localized to pre-formed microtubules (Alexa568-labeled, GMPCPP stabilized) and spun down onto coverslips; scale bar, 3 µm. Mean normalized to maximum signal and SEM of three replicate experiments (error bars) shown. See Figure S3 for comparison to in solution TPX2-tubulin co-condensation. **(C)** GFP-TPX2 and Alexa568-labeled tubulin co-localization in cytosol treated with Nocodazole to prevent MT polymerization. See Fig. 1G for experimental set up. Images taken 10 minutes into reaction (see Movie S3). Scale bar is 3 µm and TPX2 and tubulin were [2µM]. **(D)** TIRF images and **(E)** quantification of TPX2-mediated branching MT nucleation in *Xenopus* meiotic cytosol at indicated concentrations of TPX2. Shown images were taken at 1000 seconds (indicated) Cy5-labeled tubulin (red) and mCherry-EB1 (green) highlight microtubules and growing microtubule plus ends, respectively; Scale bar, 10µm. See also Movie S4. **(F)** Rate of MT nucleation as a function of TPX2 concentration for 4 independent replicates of data shown in (D) and (E). See also table 1. Rates normalized to maximum rate within an experiment. Line of best fit shown and approximate physiologically relevant range (25-100nM) highlighted. Endogenous concentration range of TPX2 (30-100 nM) indicated (also in B).

To determine whether these observations are important for TPX2 function, we first determined the physiologically relevant concentration of TPX2 for mediating branching MT nucleation. Endogenous TPX2 was removed by immunodepletion from cytosol and mono-dispersed, recombinant TPX2 was added at various concentrations. The resulting MT nucleation kinetics of branched MT networks were measured (Figure 2D-F, Movie S4) and plotted as a function of TPX2 concentration (Figure 2E). TPX2 increases the rate of MT nucleation roughly 100-fold (Table 1) in a switch-like fashion within a concentration range of 25-100nM (Fig. 2F). Strikingly, this concentration range precisely matches the endogenous TPX2 levels, and most importantly, the phase boundary of TPX2-tubulin co-condensates, which are expected to be exclusively localized along MTs at this concentration. Collectively, these observations suggest that TPX2-tubulin co-condensation on a MT could underlie TPX2’s switch-like activation of branching MT nucleation, as well as spatially bias MT nucleation to occur exclusively from pre-existing MTs.

**Table 1.**
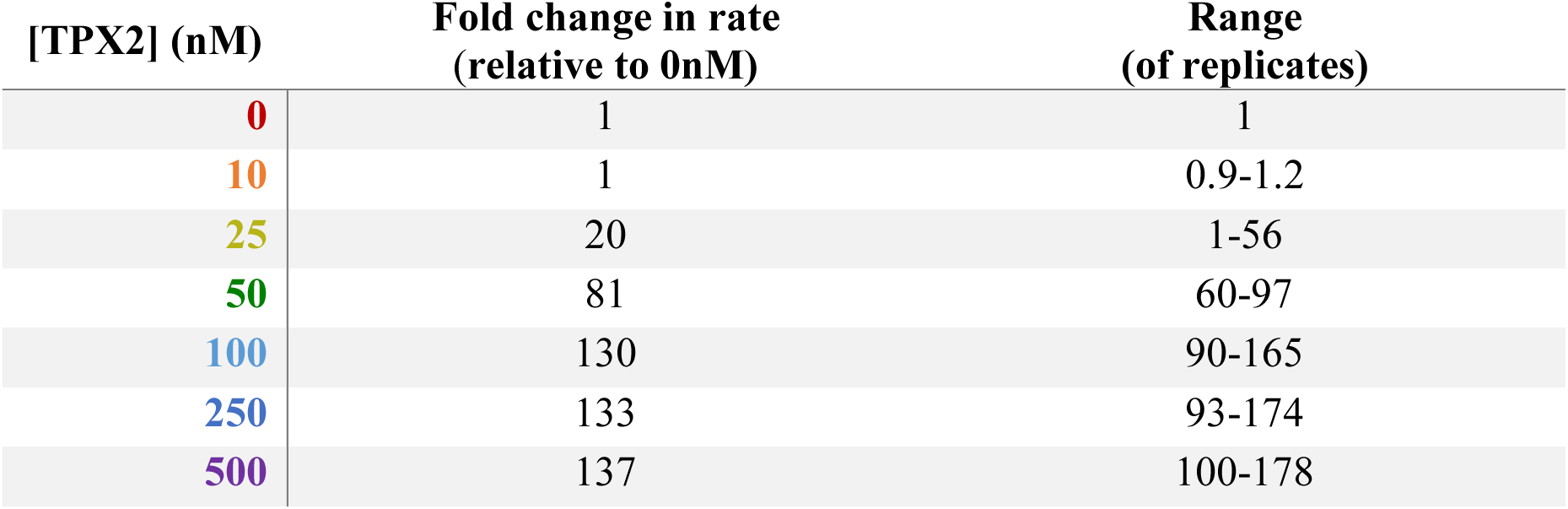
MT nucleation rates of full length TPX2. Fold change in rates of MT nucleation relative to no TPX2 added (0nM). Rates were normalized to a rate of 1 in the 0nM condition within each experimental set and these values were averaged across replicate sets (2nd column). Range of fold-change values shown of the experimental replicate sets (three or four total measurements were obtained per concertation among four replicate sets). Rates for each concentration are also displayed as individual points in Fig. 2F.

### The N-terminal region of TPX2 promotes condensation and enhances branching MT nucleation efficiency

To determine how TPX2-tubulin co-condensation contributes to branching MT nucleation, we first tested various truncations of TPX2. While co-condensation was not completely abrogated in any truncation, the disordered N-terminal constructs phase separated at much lower concentrations and had greater tubulin partition coefficients than C-terminal constructs (Fig. 3A-C, S4A-D). Although the N-terminal 1-480aa region drives the majority of TPX2 phase transition and tubulin co-condensation, it alone does not stimulate branching MT nucleation, while the minimal fragment that retains branching activity is TPX2’s C-terminal 480-716aa (Fig. 3D, S4E-F, Movie S5) (Alfaro-Aco et al., 2017). Yet, while not functional on its own, within the context of the full-length protein, the N-terminal region of TPX2 enhances the efficiency of MT nucleation about 10-fold (Fig. 3E).

**Figure 3:**
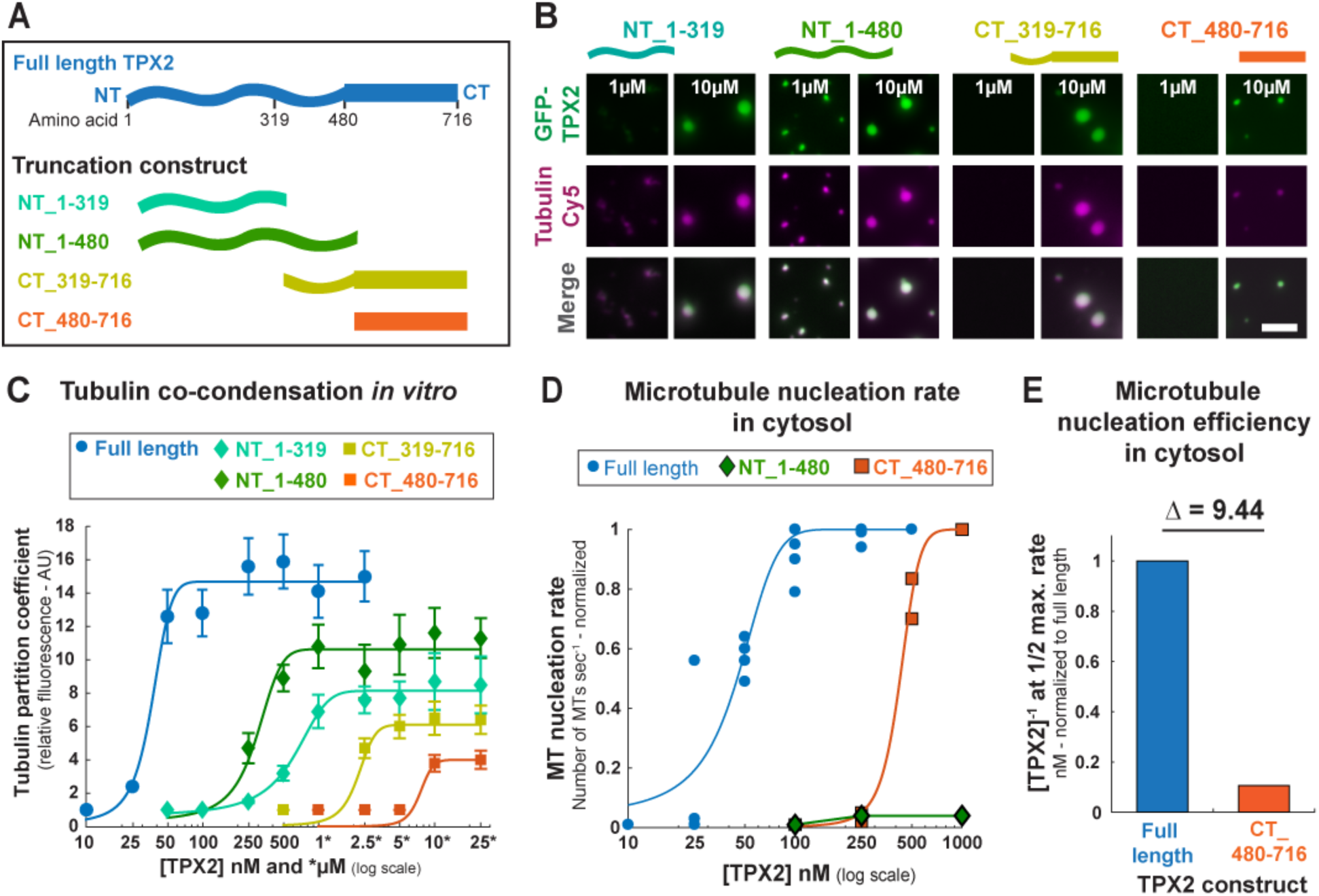
The N-terminal region of TPX2 promotes condensation and enhances branching MT nucleation efficiency. **(A)** Schematic of full-length TPX2 and various N-terminal and C-terminal truncations constructs used. **(B)** Epifluorescent images of GFP-TPX2 and tubulinCy5 co-condensates at 1µM and 10µM (equimolar concentration) for indicated TPX2 truncation construct. **(C)** Quantification of relative tubulin signal in co-condensate (partition coefficient) as a function of TPX2 concentration. Mean values with ±1 SD as error bars from a representative experiment are plotted and a line fit. See also Figure S4A-D for partition coefficient measures of GFP-TPX2 signal. **(D)** Rate of MT nucleation as a function of TPX2 concentration for indicated constructs. Full-length data previously shown (Fig. 2E). Rates for each concentration of a given construct are normalized to the maximum rate of that construct (absolute maximum rates between all constructs are in a ±2x range). Lines of best fit shown. See also Figure S4E-F for individual rate curves and movie S5. **(E)** Different efficiencies of MT nucleation for the full-length and CT_480-716 TPX2 construct. Efficiency values are the inverse of the TPX2 concentration at which half the maximum rate of MT nucleation is achieved ([TPX2]^-1^ at Rate_1/2Max_) efficiency values were normalized to full-length (efficiency of 1). The difference (fold change) in efficiencies is shown (Δ).

### TPX2-tubulin co-condensation underlies efficient branching MT nucleation

Next, we researched the mechanism by which TPX2 stimulates branching MT nucleation and the precise role of TPX2-tubulin co-condensation in this process. To do this, we replaced the N-terminal 1-480aa with various heterologous regions to generate TPX2 chimeras with distinct functionalities (Fig. 4A). We tested the ability of these chimeras to co-condense with tubulin (Fig. 4B) and promote branching MT nucleation (Fig. 4C). Due to the presence of the C-terminal minimal fragment (CT_480-716) (Alfaro-Aco et al., 2017), all of the chimeras were still capable of eliciting branching MT nucleation with reduced MT nucleation efficiency.

**Figure 4:**
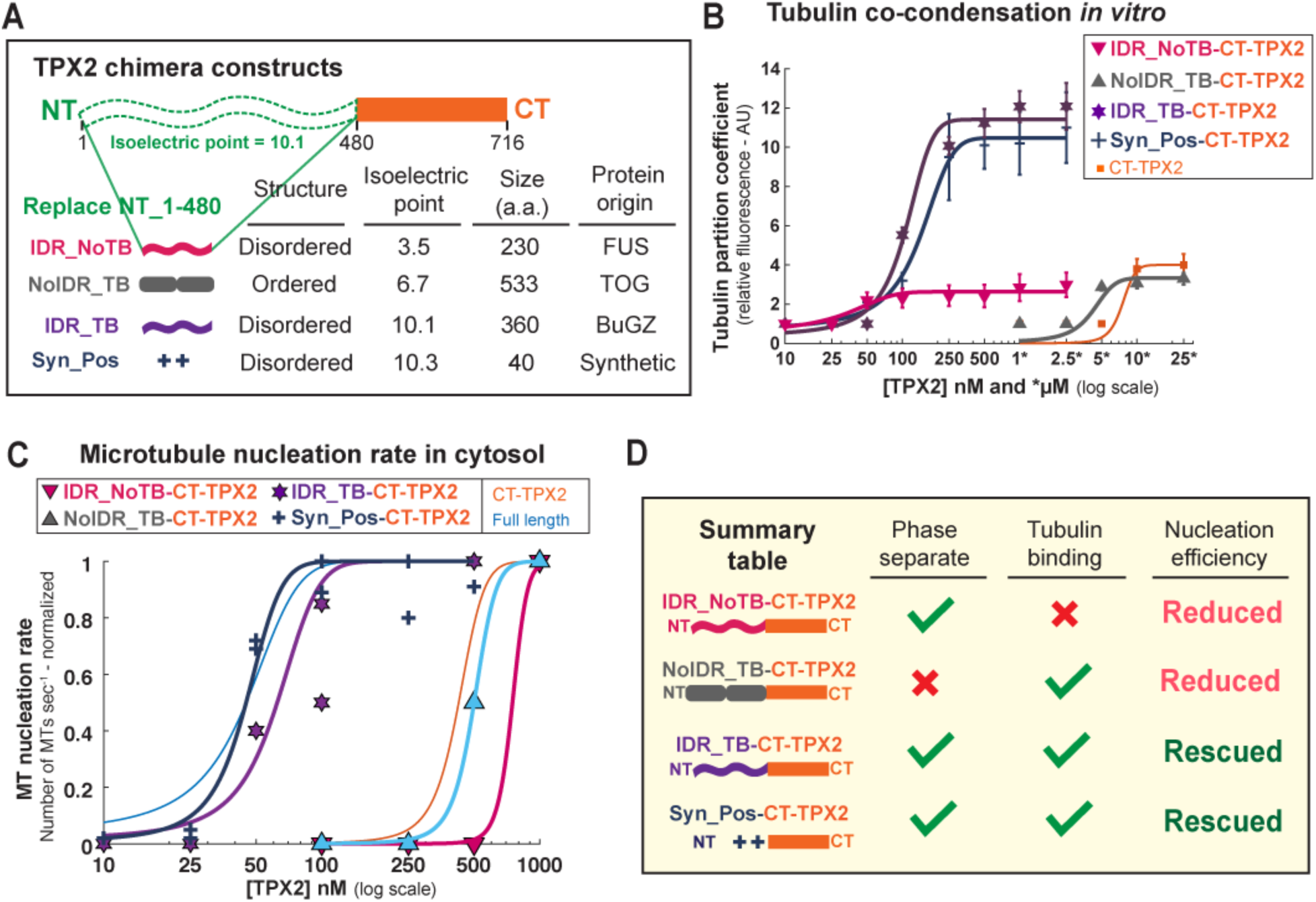
TPX2-tubulin co-condensation underlies efficient branching MT nucleation. **(A)** Schematic of chimera design: the endogenous N-terminal 1-480aa of TPX2 are replaced with exogenous domains containing distinct features shown in table. **(B)** Quantification of tubulin partition coefficient as a function of TPX2-concentration. For partition coefficient graphs mean values with ±1 SD as error bars from a representative experiment are plotted and a line fit. See also Figure S6A-B, E-F for partition coefficients of TPX2. **(C)** Rate of MT nucleation as a function of TPX2 concentration for indicated constructs. Rates for each concentration of a given construct are normalized to the maximum rate of that construct (absolute maximum rates between all constructs are in a ±2x range). Lines of best fit shown. Data for Full-length (Fig.’s 2E and 3D) and CT_TPX2 (Fig. 3D) were previously shown. See also Figure S6C-D, H-I for individual rate curves and movies S6 and7. **(D)** Summary table of results. MT nucleation efficiencies are relative to full-length TPX2.

Neither the chimera containing an N-terminal region that confers only phase separation but no additional tubulin condensation (IDR_NoTB, i.e. the intrinsically disordered region (IDR) of FUS), nor one that only associates with tubulin via two ordered TOG domains but does not phase separate (NoIDR_TB, i.e. TOG domains 1 and 2 from XMAP215), changed the reduced MT nucleation efficiency of CT-TPX2 (Fig. 4B-D, S5A-D, Movie S6). Next, the intrinsically disordered region of the MT binding protein BugZ was used. This BugZ domain phase separates and has a similar pI to the endogenous TPX2 N-terminus (IDR_TB), but does not independently promote branching MT nucleation (Fig. S5E). Remarkably, this chimera recapitulated the TPX2-tubulin co-condensation property of full length TPX2 (Fig. 4B, S5F) and, most importantly, rescued the ∼10-fold loss in MT nucleation efficiency in cytosol that arises without the endogenous N-terminal 1-480aa region (Fig. 4C-D, S6G, Movie S7).

Within TPX2, both aromatic and charged residues are highly conserved (Fig. S6A). Aromatic residues have been in implicated in tubulin condensation of the microtubule associated protein BuGZ (Jiang et al., 2015), however their removal from TPX2 does not alter tubulin co-condensation (Fig. S6B-D). We hypothesized that the abundant and conserved charged residues may mediate tubulin co-condensation and efficiency of branching MT nucleation. To directly test this, we created a chimera using a synthetic peptide (Syn_Pos) that replicated both the positive charge and intrinsic disorder of the endogenous TPX2 region (Fig. 4A). Again, this chimera fully rescued TPX2-tubulin co-condensation and the MT nucleation efficiency of full-length TPX2 (Fig. 4B-C, S6E-F). Critically, the gain-of-function chimeras (IDR_TB-CT-TPX2 and Syn_Pos-CT-TPX2) specifically discern tubulin co-condensation as the underlying property that enables the physiological role of TPX2, namely, to mediate branching MT nucleation in its endogenous range (Fig. 4D).

### Importins α/β inhibit TPX2 condensation and activity

Given the importance of TPX2-TB co-condensation to branching MT nucleation, we next sought to determine if the property was functionally regulated. The importin-α/β heterodimer inhibits TPX2 until it is released by RanGTP at the onset of mitosis (Clarke and Zhang, 2008). Because RanGTP exists as a gradient emanating from chromosomes, the effective concentration of importins-α/β is low near chromosomes but increases further away into the spindle (Clarke and Zhang, 2008). Correspondingly, we find that importins-α/β reduce both TPX2-tubulin co-condensation *in vitro* and TPX2-mediated MT nucleation in cytosol in a concentration-dependent manner: at 2-fold excess importins-α/β and higher, both are abrogated (Fig. 5A-E, Movie S8). It was previously determined that the gradient of active importins-α/β results in a sharp boundary of MT-nucleation a certain distance away from chromosomes (Kaye et al., 2018), and this distance can be modulated by altering global levels of TPX2 (Oh et al., 2016). Our findings indicate that this sharp MT-nucleation boundary could be due to TPX2-tubulin co-condensation and its threshold regulation by importins-α/β.

**Figure 5:**
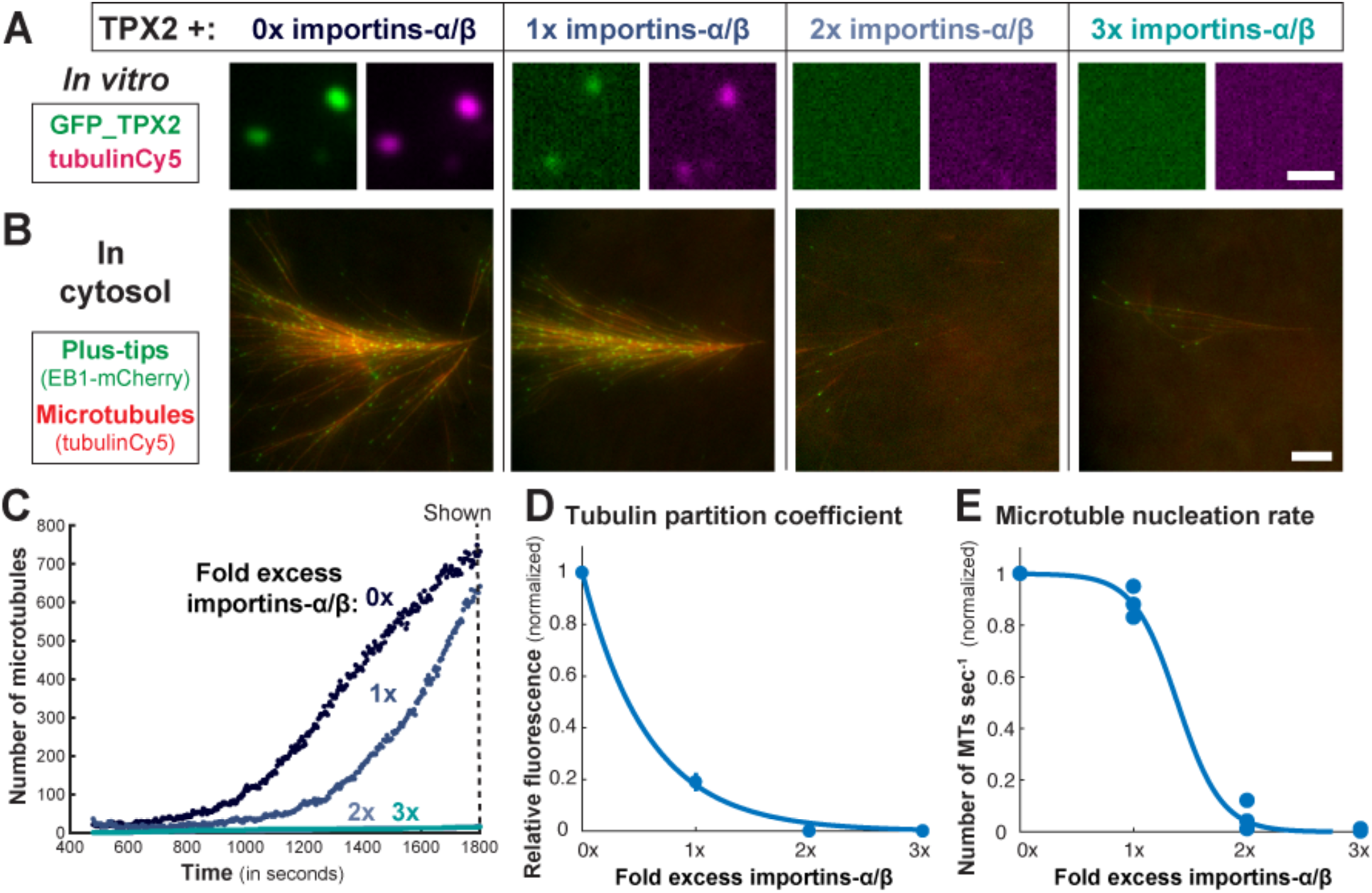
Importins α/β inhibit TPX2 condensation and activity. (**A**) Epifluorescent images of TPX2-tubulin co-condensates *in vitro* prepared with importins-α/β at indicated excess (0x = no importins-α/β) TPX2 and tubulin both at 500 nM. Scale bar, 1µm. (B) TIRF Images of TPX2-mediated MT nucleation in *Xenopus* meiotic cytosol with TPX2 and importins-α/β added at 100nM TPX2 and indicated excess of importins-α/β. Cy5-labeled tubulin (red) and mCherry-EB1 (green) highlight microtubules and growing microtubule plus ends, respectively. Images taken at 1800 Seconds. Scale bar, 10µm. See Movie S8. **(C)** Quantification of data in (B). **(D)** Quantification of relative tubulin signal (partition coefficient) as a function of excess importins-α/β. Mean values with ±1 SD as error bars shown. **(E)** Rate of MT nucleation as a function of excess importins-α/β, normalized to 0x importins-α/β, Data pooled from three experimental replicates of (B and C). Line of best fit shown.

## Discussion

Our work elucidates how TPX2 and tubulin interact on a pre-existing MT, which provides a potential mechanism for TPX2’s ability to specifically stimulate branching MT nucleation. More broadly, our findings serve as a proof-of-concept that phase separation can spatially coordinate and enhance reactions in a physiological context.

TPX2 and tubulin interact via LLPS driven by electrostatic residues located primarily in the N-terminal 1-480aa of TPX2. TPX2-tubulin co-condensates preferentially form on pre-existing MTs and at a threshold concentration of TPX2 (∼50nM) that corresponds to its endogenous and physiologically relevant concentration. This suggests that TPX2 pools tubulin via phase separation to create a local reservoir along the length of a MT (Fig. 6, right box) and that this is necessary to stimulate branching MT nucleation in an all-or-none fashion (Fig. 6, graph). Importantly, branching MT nucleation requires the C-terminal 480-716aa of TPX2 which may function through an interaction with γ-TuRC (Alfaro-Aco et al., 2017; Scrofani et al., 2015) and its co-nucleation factor XMAP215 (Flor-Parra et al., 2018; Gunzelmann et al., 2018; Roostalu et al., 2015; Thawani et al., 2018; Woodruff et al., 2017). Moreover, branching MT nucleation requires the augmin complex, which directly localizes γ-TuRC to pre-existing MTs (Song et al., 2018). Thus, while multiple proteins are essential for branching MT nucleation, our results suggest a critical role for TPX2 condensation.

**Figure 6:**
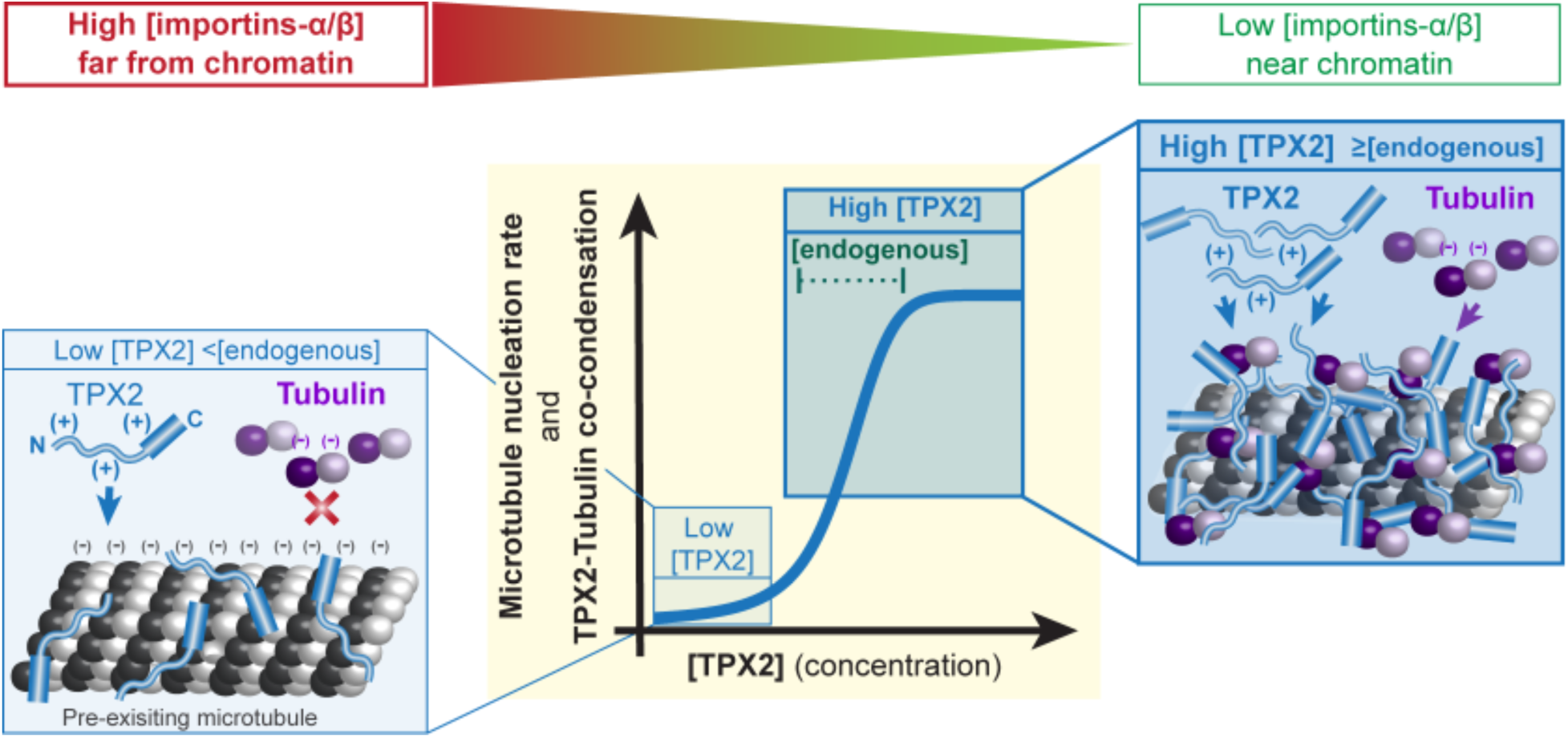
Model. Left side cartoon: At low concentrations (<endogenous) TPX2 localizes to MTs but does not recruit soluble tubulin, likely due to electrostatic repulsion (denoted by ‘+’ and ‘-’). Right side cartoon: At high concentrations of TPX2 (≥ endogenous) TPX2 co-localizes with soluble tubulin on microtubules. Center: graphical abstraction of data demonstrating that TPX2 promotes branching MT nucleation (in cytosol) and forms a co-condensate with tubulin (*in vitro*) in a switch-like fashion at or above its endogenous concentration. Top gradient: relative importin-α/β levels existing as a gradient around chromosomes also affect TPX2-tubulin co-condensation and TPX2-mediated branching MT nucleation.

Loss of TPX2 results in reduction of spindle microtubule mass and mitotic delay (Neumayer et al., 2014), which is early embryonic lethal in mice (Aguirre-Portolés et al., 2012) and causes apoptosis in proliferative somatic cells (Neumann et al., 2010). TPX2 overexpression leads to aberrant additional puncta of MT nucleation (Brunet et al., 2008; Ma et al., 2010) that our data suggests correspond to TPX2-tubulin condensates. Furthermore, TPX2 overexpression is observed in 27% of all cancer types (Neumayer et al., 2014) and the degree of TPX2 overexpression tightly correlates with disease lethality (Carter et al., 2006; Wang et al., 2018). In sum, maintaining appropriate TPX2 levels is essential for healthy cell division. Our work provides a potential mechanistic framework to understand the relationship between TPX2 levels and its associated microtubule nucleation activity, which could educate future therapeutic efforts.

Almost two decades ago, TPX2 was identified as the major downstream factor of RanGTP that is required for MT generation from chromatin, but the molecular mechanism of its regulation has remained unclear (Gruss et al., 2001; Neumayer et al., 2014). Our observation of complete inhibition of both TPX2-tubulin co-condensation and TPX2-mediated MT nucleation by importins-α/β could explain how TPX2 coordinates a sharp boundary of nucleation a certain distance from chromatin within the spindle (Fig. 6, above graph) (Kaye et al., 2018; Oh et al., 2016). Phase separation as a means to convert gradient signals into binary responses has implications for many biological processes including signaling and morphogenesis (Csizmok et al., 2016). Collectively our findings suggest that TPX2-tubulin co-condensation is a regulated process that may underlie the all-or-nothing activation of branching MT nucleation.

It has been proposed that phase separated condensates act as reaction crucibles to enhance rates and efficiency (Banani et al., 2017; Castellana et al., 2014; Shin and Brangwynne, 2017). This conclusion is based on many *in vitro* studies (Hernández-Vega et al., 2017; Jiang et al., 2015; Li et al., 2012; Strulson et al., 2012; Su et al., 2016; Woodruff et al., 2017), but remains poorly understood in a physiological context (Boija et al., 2018; Du and Chen, 2018; Zhao et al., 2015). Our results demonstrate that phase separation of TPX2 and tubulin improves the efficiency of branching MT nucleation in cytosol at least 10-fold. The fact that TPX2 selectively phase separates onto a pre-existing MT enables the autocatalytic amplification of MTs via branching MT nucleation, which increases MT nucleation rates up to 100-fold. In summary, we established that phase separation of a MT nucleation effector can enhance both reaction efficiency and rate whilst also spatially coordinating activity. Phase separation mediated mechanisms are likely at play in a broad set of cellular processes and this will be important to explore in physiological contexts.

## Materials and Methods

*Contact for Reagent and Resource Sharing*

Further information and requests for resources should be directed to and will be fulfilled by lead contact Dr. Sabine Petry

### Protein constructs, expression and purification

#### E. Coli Strains

DH5α cells (New England Biolabs (NEB): C2987I) were used for all subcloning steps. Rosseta2 cells (Fisher: 71-403-4) were used to express proteins for purification. Cells were grown at various volumes in LB Broth (Sigma: L3522) prepared according to supplier’s instructions.

#### Protein constructs

BSA (Fisher: 23209) and Tubulin (PurSolutions LLC: 032005) are bovine versions and proteins were acquired directly from vendors. FUS and Ran are the human versions. All remaining proteins are *Xenopus laevis* versions. DNA sequences were sourced from in-house plasmids, the *Xenopus laevis* Gene Collection (Source Biosciences), synthesized (GenScript, Sigma) or were gifted (FUS-IDR plasmid was a gift from Cliff Brangwynne).

All TXP2 constructs were cloned as N-terminally tagged Strep6xHisGFP-TEV-TPX2 fusions using a modified pST50 vector (Tan et al., 2005) and cloned via Gibson assembly (NEB: E2611L). An identical strategy was used to generate StrHisTEV-mCherry-FL_TPX2, StrHisBFP-TEV-IDR_TB, StrHisBFP-TEV-RanQ69L, EB1-mCherry6xHis *(27)*, GST-importin α, and GST-importin β. Insert fragments were PCR amplified from plasmids containing the indicated gene which was unmodified from its wild type sequence. Exceptions are: Ran, which has a single site mutation Q69L to render it dominant positive, and NTΔYF_TPX2 and Syn_Pos-CT_TPX2 which were both custom synthesized (Genscript (NTΔYF) and Sigma (Syn_Pos): personal quote). The Syn_Pos (synthetic positive) protein sequence is: AKKRKAGDSEGSEGAKKRKAAKKRKAGDSEGSEGAKKRKA. This sequence was generated by interspacing nuclear localization-like sequences (e.g. KKRK) with inert a.a.’s G, S, A, as well as, oppositely charged E. The construct was designed to have a similar theoretical isoelectric point (pI) to the endogenous NT1-480aa of TPX2. All constructs were fully sequenced and confirmed to have no errors.

#### Expression and purification

All constructs were transformed into Rosseta2 *E. coli* cells and were grown in temperature-controlled incubators shaking at 200RPM. For protein expression, cells were grown at 37°C (0.5-0.7 OD_600_), then cooled to 16°C and induced with 0.75 mM isopropyl-β–D-thiogalactoside (IPTG) for another 7 hours at 27°C. Cell pellets were collected and flash frozen for future protein purification.

For all TPX2 constructs, cells were lysed using an EmulsiFlex (Avestin) in lysis buffer (0.05M Tris-HCl, 0.015M Imidazole, 0.75M NaCl, pH 7.75) containing 0.0002M phenylmethylsulfonyl fluoride (PMSF), 0.006M β-mercaptoethanol (βME), cOmplete™ EDTA-free Protease Inhibitor tablet (Sigma 5056489001), and 1000U DNase I (Sigma 04716728001). Lysate was centrifuged in a F21-8×50y rotor using a Sorvall RC6+ at 17,000RPM for 25min. Clarified lysate was bound to pre-washed Ni-NTA agarose beads (Qiagen 1018236) in a gravity column and beads were washed with 10 column volumes lysis buffer. Protein was eluted in lysis buffer containing 200mM Imidazole, and then further purified via gel filtration (Superdex 200 HiLoad 16/600, GE Healthcare – 28-9893-35) in CSF-XB buffer (0.01M Hepes, 0.002M MgCl, 0.0001M CaCl, 0.004M Ethylene glycol-bis(2-aminoethylether)-N,N,N′,N′-tetraacetic acid (EGTA), 10% w/v sucrose, pH-7.75) containing either 0.1M KCl for extract assays or 0.5M KCl for condensate assays. Peak fractions were pooled, concentrated (Amicon Ultra, ThermoFisher - various sizes), flash frozen and stored at −80°C. Untagged TPX2 was generated by cleaving Strep6xHisGFP-Tev-FL protein with TEV protease at 100:1 TPX2:TEV protease molar ratio overnight at 4°C while dialyzing into cleavage buffer (0.02M NaPO_4_, 0.5M NaCl, 0.006M βME, and 0.0002M PMSF, pH −7.5). Cleaved TPX2 was collected as the flow-through of the reaction mixture added to NiNTA agarose beads.

For GST-importin α and GST-importin β, clarified lysates were prepared in the same way with the exception of the lysis buffer (0.05M Tris-HCl, 0.138M NaCl, 0.0027M KCl pH 8) and they were bound to a GST affinity column (GSTrap™ Fast Flow, GE Healthcare: 17-5131-02). The column was washed (0.02M NaPO_4_, 0.15M NaCl, 0.006M βME, and 0.0002M PMSF, pH - 7.5), protein was eluted (0.02M Tris-HCl, 0.15M NaCl, 0.01M L-Glutathione, 0.006M βME, and 0.0002M PMSF, pH – 7.5) and peak fractions were pooled. GST-importin β was cleaved to remove GST and purified in the same manner as untagged TPX2. Strep6xHisBFP-RanQ69 and EB1-6xHismCherry were purified as described previously (Alfaro-Aco et al., 2017; Thawani et al., 2018). Pure tubulin and BSA were labeled with commercial NHS-conjugated dyes (Cy5 or Alexa568) according to supplier’s instructions (Sigma: GEPA150101). A similar method was used to conjugate biotin to tubulin (EZ-Link™ NHS-PEG4-Biotin, ThermoFisher: 21329). Conjugated tubulin was further purified for MT competent tubulin by a series of MT polymerization and pelleting rounds. The percentage of labeling was ≥ 60% for all purifications. In direct comparisons of fluorescently conjugated proteins (i.e. BSACy5 and tubulinCy5) batches were used that had matched percentage of labeling.

All proteins were flash frozen in CSF-XB buffer at either 0.1M of 0.5M KCl and stored at −80°C. Before use, all proteins were pre-cleared of aggregates via ultracentrifugation at 80,000 RPM for 15min in a TLA100 rotor in an Optima MAX-XP ultracentrifuge at 4°C. Protein concentrations were determined by Coomassie-blue densitometry measures of a concentration series of the protein of interest and a BSA standard run on the same SDS-PAGE gel.

### Image collection and processing

The imaging technique used is indicated in each corresponding figure legend. Total Internal Reflection Fluorescence (TIRF), Epifluorescence (Epi), and Differential Interference Contrast (DIC) microscopy methods were carried out on a Nikon TiE microscope with a 100X, 1.49NA oil immersion objective and an Andor Zyla sCMOS camera. Confocal microscopy was carried out on a Nikon TiE microscope with a 63X, 1.45NA oil immersion objective, a Yokogawa CSU 21 disk module and a Hamamatsu ImageEM EM-CCD camera. Fluorescence recovery after photobleaching (FRAP) was carried out on a Nikon A1 point scanning microscope using a 63X, 1.45NA oil immersion objective. All experiments using *Xenopus* egg cytosol were carried out at 18°C in a temperature-controlled room.

NIS-Elements software was used for all image acquisition. All images within a data set were taken with identical imaging parameters. Binning was not used except in the case of extract branching MT nucleation assays. FIJI and MATLAB were used for all image analysis. Images within a figure panel were processed with matched brightness and contrast windows for each color to allow direct comparison of intensities. In a few cases, enhanced contrast was used to emphasize non-association events, in which case it was indicated in the legend. These cases include figures 2A, S1D, S3D and F, which have enhanced contrast as the main image only (2A and S3F) or have an enhanced contrast image in addition to the matched contrast image (S1D and S3). All images are representative crops. Subtraction of background signal was not used, except in the case of GFP_TPX2 signal in Fig. 1I. Oblique TIRF was used in these and other cases (indicated in figure legends) to visualize GFP-TPX2 signal, which cannot be seen in TIRF due to high levels of TPX2 bound to the coverslip.

### Condensate (phase separation) assays

#### Standard assay

Proteins were diluted to 5x final concentration in a CSF-XB Buffer containing 500 mM KCl salt, then diluted 1:4 in CSF-XB containing no salt at 4°C to reach a final salt concentration of 100mM. The reaction mixture was immediately pipetted into a 5uL volume coverslip flow chamber (constructed with double-stick tape, a glass slide and a 22×22mm coverslip). The slide was placed coverslip-side down into a humidity chamber for 20 minutes at room temperature to allow condensates to settle; reaction was then imaged. Multiple mounting strategies were explored, all yielding similar results. For condensates containing more than one protein, the molar ratio was always 1:1, unless otherwise indicated. Crowding agents were never used.

#### Partition coefficient measurements

Partition coefficient is defined as the difference in mean intensity of a condensate compared to background, i.e. apparent relative enrichment. The ‘Color Threshold’ (Otsu thresholding) and ‘Particle Analyzer’ functions on FIJI were used to identify and quantify condensate intensity, respectively. For each concentration, construct, and condition at least 100 condensates were analyzed. All statistical analysis and graph generation was carried out in MATLAB.

#### Live fall-down assay

A coverslip-bottomed CultureWell (Grace BioLabs: 112359) containing CSF-XB buffer with no salt was positioned and focused on a microscope and acquisition was started. 10x concentration of protein was diluted directly into the well to achieve a 1:9 dilution and a final salt concentration of 100mM. Time 00:00 (minutes:seconds) corresponds to the addition of the protein.

#### FRAP (Fluorescence Recovery After Photobleaching)

Condensates were prepared as in the live fall-down assay and allowed to settle for 5 minutes. Focus was set just above the coverslip and three Regions Of Interest (ROIs) of equal size and geometry were placed (1) within the condensate to be photobleached, (2) the background, and (3) within a nearby condensate. Photobleaching was carried out and the intensity of each ROI recorded every second for the first 10 seconds and every 10 seconds thereafter over a 10- to 20-minute acquisition. Recovery in the photobleached ROI was normalized to any changes in intensity in the background and nearby condensate ROI due to global bleaching. Global photobleaching in excess of 5% of the starting intensity was never observed. If the intensity in the nearby condensate ROI changed, that FRAP acquisition was discarded.

#### *In vitro* MT Polymerization assay

Coverslip flow chamber was first blocked with κ-casein (Sigma: C0406-1G), and then coated with PEG-biotin (Rapp Polymere: 133000-25-20) and NeutrAvidin™ (ThermoFisher: A2666), each in BRB80 buffer (0.08 M Pipes, 0.001 M MgCl_2_, 0.001 M EGTA, pH– 6.8), and with a BRB80 wash in between each step. Condensates containing tubulin and TPX2 were prepared as in the standard assay but in this case diluted into BRB80 buffer containing oxygen scavengers and 1 mM GTP for final salt concentration of 50 mM KCl and 80mM Pipes. Final concentrations were 10 µM bovine tubulin with 10% labeled Cy5-tubulin, 2% biotinylated-tubulin and 1 µM GFP-TPX2. Condensate mixture was pipetted into the coverslip flow chamber and imaged.

#### Measure of TPX2 soluble pool (i.e. phase boundary)

Condensates were prepared as in the standard assay at concentrations indicated and gel samples were prepared as inputs. Samples were spun at 80,000 RPM for 15min in a TLA100 rotor in an Optima MAX-XP Ultracentrifuge and gel samples were prepared from supernatants (soluble pool). Soluble pool sample concentrations were obtained via densitometry measurements of silver-stained SDS-PAGE.

#### Phase boundary as a function of [salt] and [TPX2]

Condensates were prepared as in the standard assay at protein and salt concentrations indicated. A particular [salt]:[TPX2] condition was scored to have phase separated condensates if the average signal in apparent condensates was at least four times higher than background (i.e. partition coefficient ≥4). Conditions were scored to not contain condensates if they fell below this cutoff (only occurred in a three [salt]:[TPX2] conditions) or if no apparent condensates were observed (majority of cases).

### Microtubule localization assays

#### Concentration Series

50µL of a dilute solution of GMPCPP-stabilized microtubules containing 10% Alexa568-labled tubulin were mixed with a 50µL solution of monodispersed TPX2 (GFP-tagged) and tubulin (Cy5Labled), both at equal molar concentrations in 0.1M KCl CSF-XB buffer containing 1mM GTP. Mixture was immediately spun onto Poly-L-Lysine (Sigma: P8920) coated round coverslips through a 20% glycerol cushion at 12000 RMP in an HB-6 swinging bucket rotor using a Sorvall RC 6+ centrifuge. Coverslips were mounted with ProLong® Diamond Antifade Mountant (ThermoFisher: P36970), which was allowed to cure before imaging.

#### Localization of tubulin to MTs

Sample was prepared in the same way as the concentration series experiment. Protein mixtures contained equal molar concentrations of monodispersed proteins (TPX2, tubulin, BSA, and/or GFP, as indicated in figure legend).

#### Time Series of TPX2 localization

80µL of a dilute solution of GMPCPP-stabilized microtubules containing 10% Alexa568-labled tubulin and 10% biotinylated-tubulin were attached via anti-biotin antibodies adhered to the surface of a blocked (κ-casein) coverslip-bottomed CultureWell (Grace BioLabs: 112359). The microscope stage was positioned and focused on MTs, after which acquisition started. 10x concentration of mono-dispersed TPX2 was diluted directly into the well containing BRB80 buffer with 1mM GTP and oxygen scavengers to achieve 1X TPX2 concentration and a final salt concentration of 50mM KCl and 80mM Pipes. Time 00:00 corresponds the frame before any TPX2 signal is observed (approximately 5 seconds after the addition of protein).

### *Xenopus* egg cytosol assays

#### Cytosol preparation

Cytosol naturally arrested in meiosis II was prepared as described in (Hannak and Heald, 2006). Briefly, stage VI (mature, meiotically arrested) *Xenopus laevis* eggs were collected after an overnight laying period. Eggs from individual frogs were kept separate but prepared in parallel, and typically 2-3 batches of eggs were used. Egg jelly coats were removed and cytosol was fractionated away from egg yolk, membranes, and organelles by centrifugation (10200 RPM in HB-6 for 15 minutes). Eggs were constantly maintained at 18°C via preparation in a temperature-controlled room. Undiluted cytosol was collected, supplemented with Cytochalasin-D, protease inhibitors, ATP, and creatine phosphate and kept at 4°C until use. All reactions shown within a single image panel were acquired from the same cytosol preparation imaged in a single session.

#### Extract overlaid onto condensates

1 µL of TPX2 condensates, prepared as described in the standard assay, were pipetted onto the center of an untreated coverslip and overlaid with 5 µL *Xenopus* cytosol (containing mono-dispersed Alexa568-labled tubulin). Extract overlay was carried out either immediately or after condensates were aged for the indicated amount of time (Fig. S2D-F). A slide was gently placed on top of the mixture, which marked the start of the reaction (time 0 sec). The sample was imaged via oblique-TIRF microscopy. Condensate and branched MT network mass in Figure S2B were calculated by creating fixed size convex hulls around each and conducting integrated intensity measurement across all frames.

#### Immunodepletion

Preparation of *Xenopus* cytosol immunodepleted of TPX2 was prepared as described in (King and Petry, 2016). Briefly, immunoaffinity purified antibodies against TPX2 or an unspecific IgG control antibodies were conjugated to magnetic Dynabeads Protein A (ThermoFisher: 1002D) at 4°C overnight. Antibody-conjugated beads were split into two equal volume aliquots; supernatant was removed from one aliquot using a magnetic block and *Xenopus* cytosol was added. Beads were gently suspended in cytosol every 10 minutes for 40 minutes. Cytosol was removed from beads (using magnetic block) and then subjected to another round of depletion using the same procedure with the second aliquot of antibody-conjugated beads. Immunodepletion was assessed via functional assays.

#### Branching MT nucleation assay

MT nucleation reactions were carried out as described in (King and Petry, 2016). Briefly, *Xenopus* cytosol was supplemented with fluorescently labeled tubulin ([1 µM] final) to visualize MTs, mCherry-fused End Binding protein 1 (EB1) ([100 nM] final) to track MT plus ends, and sodium orthovanadate ([0.5 µM] final) to inhibit motors and prevent MT gliding. Mono-dispersed purified TPX2 constructs (and other proteins) were added at specified concentrations. In the case of + nocodazole (Fig 1G), it was added at a final concentration of 0.3mM. In experiments using immunodepleted cytosol (Figures 2 and 3) a constitutively active form of RanGTP (Ran^Q69L^) ([7.5µM] final) was added to prevent sequestration of TPX2 by endogenous importins. In all experiments 0.1 M KCl CSF-XB buffer was used to match total dilution (25% of extract volume across all experiments). The reaction mixture was prepared on ice, then pipetted into a coverslip flow chamber at 18°C, which marked the start of the reaction. Reaction was imaged via TIRF microscopy for 20-40minutes. See ‘Analysis of branching MT nucleation’ section for details of time-lapse acquisitions. All reagents used were in 0.1 M KCl CSF-XB buffer.

#### Analysis of branching MT nucleation

In all cases, at least three replicates of the concentration series for each construct was carried out. In these replicates, concentrations were tested in parallel on a single slide set-up with multiple flow chambers and imaged at time intervals of usually 30sec to 5min intervals over the course of 30 minutes. These intervals are longer than the 2 or 4 second time intervals that are shown in the main figures. In these replicate experiments, multiple constructs or concentrations could be assessed in parallel using a single cytosol prep, which has a finite lifetime (∼several hours). These replicates served to verify the patterns observed in the time-lapse movies required for MT nucleation rate analysis - described below.

A custom MATLAB software (Thawani et al., 2018) was used to measure the number of MTs over time. EB1 signal on MT plus ends in the entire field of view was used to determine MT number within a single frame. EB1 detection was achieved via the plus-end tracking module of uTrack (Applegate et al., 2011) applied to the entire movie (typically 400-750 frames, 2 or 4 seconds/frame). Parameters were optimized for each movie according to visual assessment of tracking accuracy. MT nucleation curves were generated by plotting the number of EB1 detections per frame over time. Branching MT nucleation rate is defined as the slope of the linear portion of the MT nucleation curve– i.e. the initial lag and eventually saturation are not used. In instances where branching MT nucleation was not observed (i.e. constant rate of de novo nucleation), rate corresponds to the slope of the entire MT nucleation curve. This quantification method is consistent with previous publications (Alfaro-Aco et al., 2017; Song et al., 2018; Thawani et al., 2018).

For each construct, a number of concentrations were quantified to generate MT nucleation rates within that series. Rates were normalized to the maximum rate within the series and plotted as a function of concentration. A logistic regression equation and the curve fitting tool in MATLAB were used to derive line of best fit. In these experiments, time-lapse movies (∼30 minutes in duration) are acquired at 1 frame/2 or 4 seconds for each concentration, for each construct. Typically, 2 to 6 movies can be acquired per cytosol, given its lifetime. These movies are used to carry out MT nucleation rate analysis described above. For most constructs at least two independent concentration series and MT nucleation rate quantifications were carried out. Exceptions are IDR_NoTB-CT-TPX2 and TB_IDR-CT-TPX2, where a single concentration series (using time-lapse movies) and quantification was carried out. Note that all quantification of every construct was verified with at least three independent replicates (independent cytosol preparations), as mentioned above.

### Other analyses

#### Prediction of protein disorder and secondary structure

Disorder prediction was carried out in IUPred (Dosztányi et al., 2005) which generates a per amino acid prediction of disorder on a scale of 0 to 1. This data was exported and converted into a heatmap using the heatmap function on MATLAB. Secondary structure predictions of TPX2 were carried out previously (Alfaro-Aco et al., 2017).

#### Measure of TPX2 soluble pool (i.e. phase boundary)

Condensates were prepared as for the standard assay at concentrations indicated and gel samples were prepared as inputs. Samples were spun at 80,000 RPM for 15min in a TLA100 rotor in an Optima MAX-XP Ultracentrifuge and gel samples were prepared from supernatants (soluble pool). Soluble pool sample concentrations were obtained via densitometry measurements of silver stained SDS-PAGE.

#### Plots and graphics

All plots (graphs) were generated in MATLAB. All graphics were made in Adobe illustrator.

### Quantification and Statistical analysis

All number or replicates (n) and statistical analyses are indicated in corresponding figure legends.

#### In vitro assays

For all *in vitro* assays, at least three independent replicate experiments were conducted. Either all quantifications are pooled and displayed, or quantifications from a single representative experiment are shown, indicated in figure legend. In the latter cases (e.g. partition coefficient measurements) data presented corresponds to the images shown (if applicable).

#### *Xenopus* egg cytosol assays

All experiments using *Xenopus* egg cytosol were reproduced at least three times using separate cytosol preparations. Similar results were seen in all replicates.

### Experimental organisms Used

#### *E. Coli* Strains

DH5α cells were used for all subcloning steps.

Rosseta2 cells were used to express proteins for purification. Cells were grown at various volumes in LB Broth (Mentioned in Key resource table → Sigma: L3522) prepared according to supplier’s instructions.

#### *Xenopus leavis* frogs

Mature (3 to 7-year-old) female *Xenopus leavis* frogs laid eggs that were used for cytosolic extract experiments. Animal housing, maintenance, and egg harvesting were all carried out to highest standards and in accordance to approved IACUC protocols and guidelines.

## Supporting information

Supplemental_Movie_1_TPX2_Fusion

Supplemental_Movie_2_TPX2_cytosol

Supplemental_Movie_3_TPX2_cytosol_nocadozol

Supplemental_Movie_4_TPX2_concentration_series

Supplemental_Movie_5_TPX2_Truncations

Supplemental_Movie_6_TPX2_Chimeras_NoRescue

Supplemental_Movie_7_TPX2_Chimeras_Rescue

Supplemental_Movie_8_Importins

## Acknowledgments

We thank members of the Petry Lab for helping with this work, including Ray Alfaro-Aco, Michael Rale, Mohomad Safari, and Akanksha Thawani. We are especially grateful to Cliff Brangwynne for experimental suggestions and the Petry Lab, Ibrahim Cisse and Kassandra Ori-McKenney for feedback on the manuscript. This work was supported by Ph.D. training grant T32GM007388 by NIGMS of the National Institutes of Health (to M.R. King), as well as the New Innovator Award of NIGMS of the National Institutes of Health (DP2), the Pew Scholars Program in the Biomedical Sciences, and the David and Lucile Packard Foundation (all to S. Petry).

## Author contributions

M.R.K. and S.P. conceived the project, designed the experiments, and wrote the manuscript. M.R.K. generated and characterized reagents and tools, and performed and analyzed experiments.

## Declaration of interests

Authors declare no competing interests.

## Supplemental Information

**Fig. S1.**
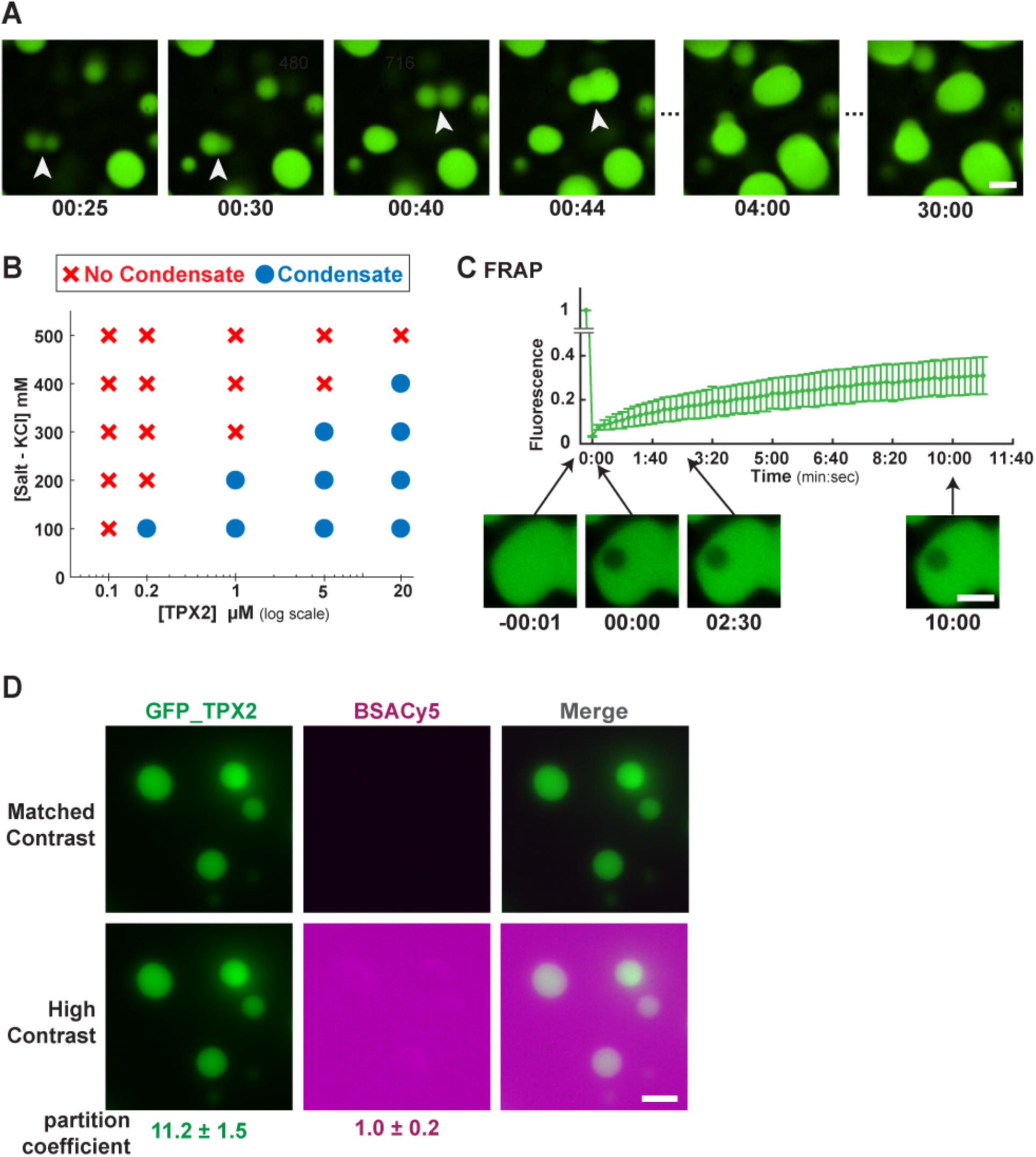
TPX2 phase separates into a liquid-like condensate. **(A)** Confocal microscopy images of GFP-TPX2 condensates falling and fusing in a coverslip-bottomed well. Select frames from time-course movie shown. 00:00 (minutes:seconds) corresponds to when GFP-TPX2 was added to the well. Arrowheads indicate fusion events. GFP-TPX2 at 20µM. Scale bar, 3µm. **(B)** Phase diagram of GFP-TPX2 at indicated salt (mM) and protein (µM) concentrations. Blue circles indicate presence and red crosses indicate absence of condensates**. (C)** Fluorescence recovery after photo-bleaching (FRAP) of mCherry-TPX2 condensates (pseudo-colored green), acquired via confocal microscopy. Mean and SEM of three replicate experiments (error bars) shown. Example images shown immediately before (−00:01) and after photobleaching (00:01). Also shown are two time-points into recovery. Scale bar, 3µm. **(D)** Epifluorescent image of GFP-TPX2 condensates prepared with Cy5-labeled BSA, both at 4 µM. Scale bar, 3µm. In upper panel the contrast is matched to main figure 1C, and enhanced in the lower panel to illustrate the absence of BSA enrichment. Partition coefficient value is the mean with ±1 standard deviation (SD) computed from at least 100 condensates in an experimental set.

**Fig. S2.**
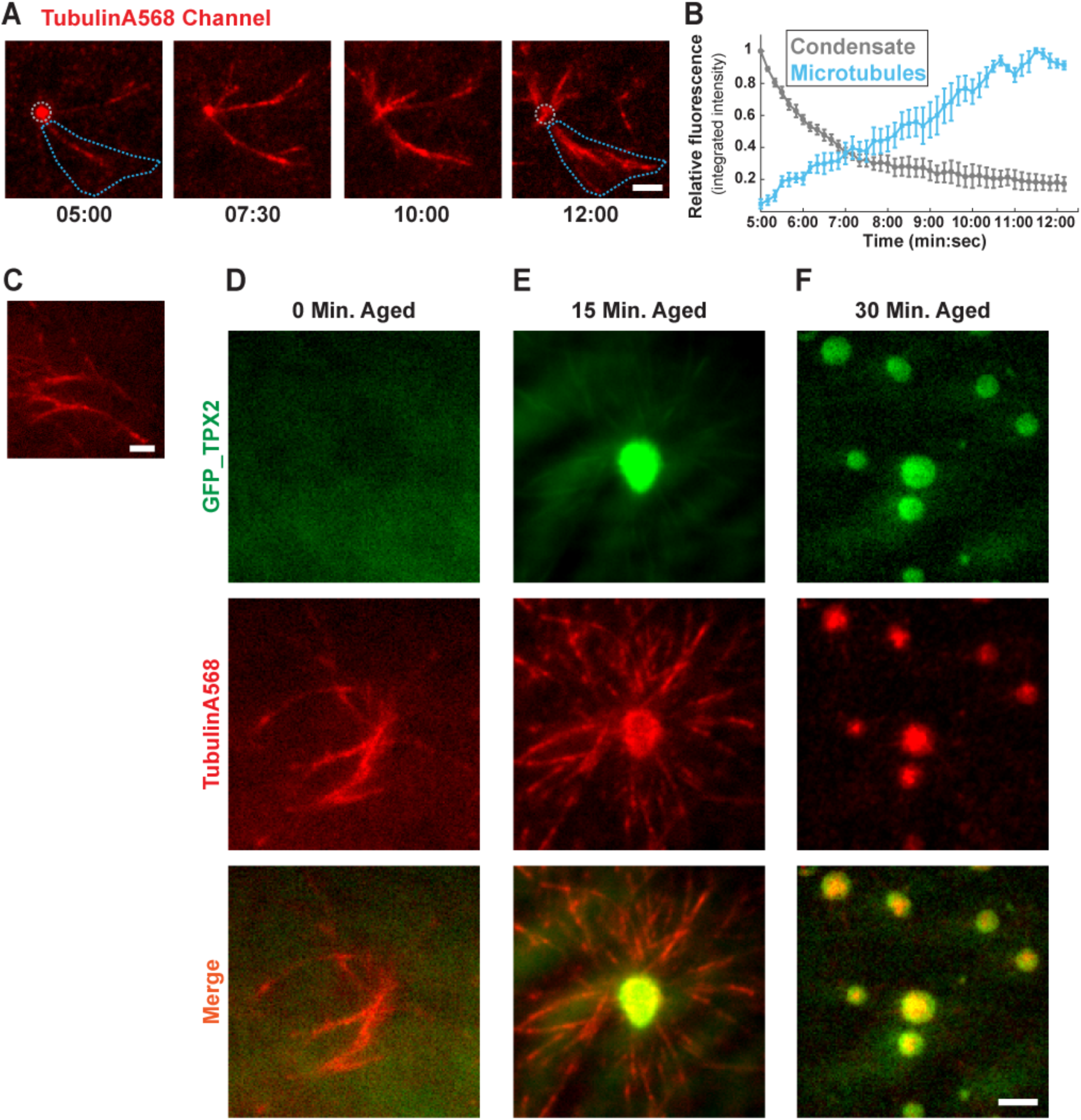
TPX2 condensate age affects MT nucleation. **(A)** In the same experiment shown in Fig. 1I, the tubulin channel imaged over time (minutes:seconds) and depicted. **(B)** Quantification of integrated tubulin signal from indicated areas corresponding to initial condensates (grey) and MT fan structures (blue). Mean values shown as circles with ±1 SD shown as error bars from five separate analyses. **(C)** An alternate field from same experiment as Fig. 1I and Sup. Fig. 2A at 15 minutes into the reaction. Scale bar, 3µm. GFP-TPX2 condensates aged for **(D)** 0 minutes **(E)** 15 minutes and **(F)** 30 minutes were overlaid with cytosol containing mono-dispersed Alexa568-labeled tubulin (see schematic in Fig. 1H). Images acquired via oblique TIRF microscopy 20 minutes after sample preparation. GFP-TPX2 and tubulin (Alexa568-labeled) channels and merge shown. TPX2 at 2 µM; scale bar, 3µm.

**Fig. S3.**
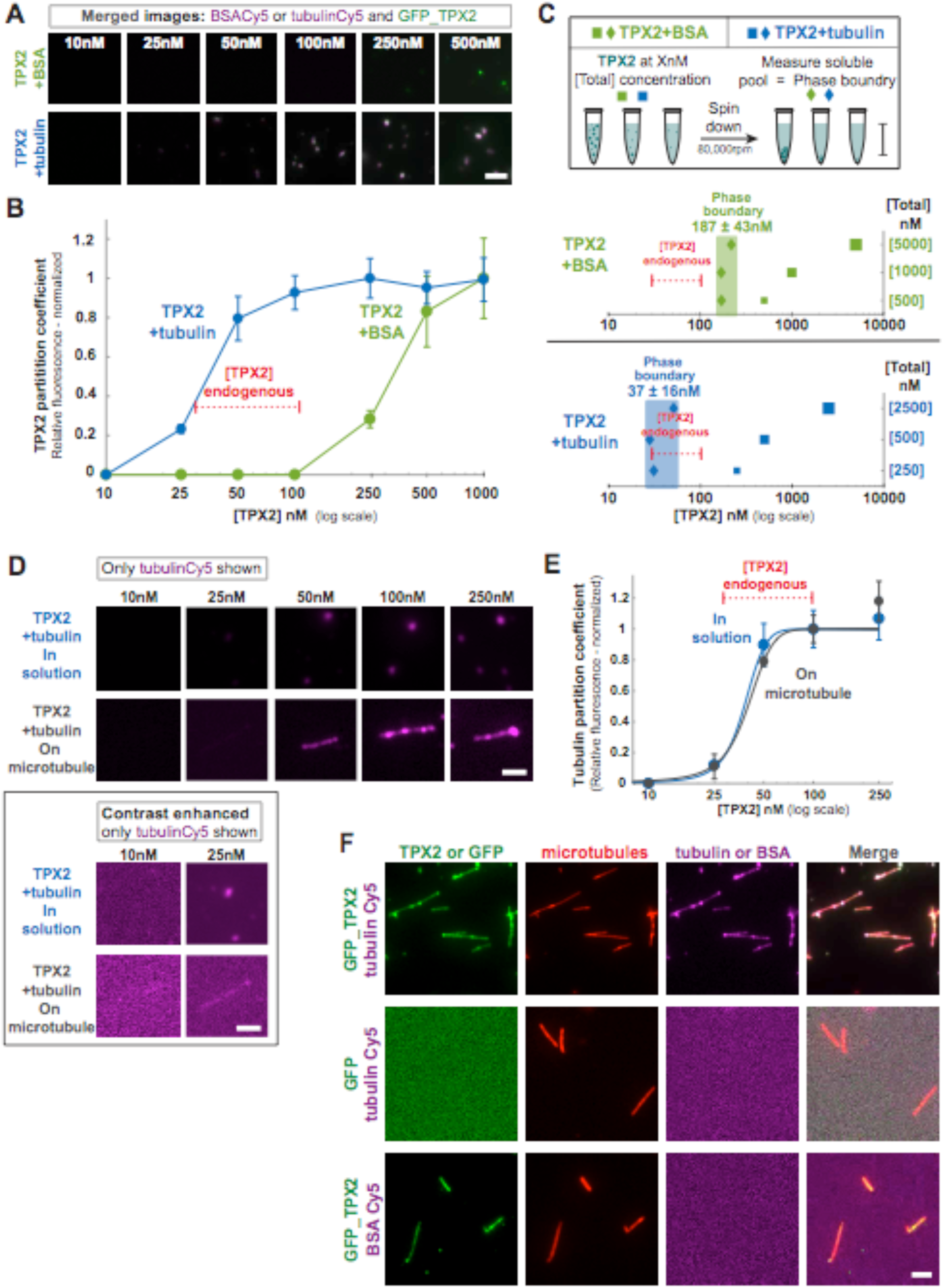
The phase boundary of TPX2 is lowered by tubulin, and TPX2-tubulin co-condensates specifically form on microtubules. **(A)** Epifluorescent images of TPX2 condensates with BSA and TPX2-tubulin co-condensates *in vitro*. BSA, tubulin, and TPX2 are equimolar at indicated concentrations. Scale bar, 2µm. **(B)** Graph of partition coefficient of GFP-TPX2 normalized to the maximum partition coefficient among the concentrations shown. Mean values shown as circles with ±1 SD shown as error bars for condensates formed in the presence of BSA (green) or tubulin (blue) and plotted as a function of concentration. At least 100 condensates per concentration were analyzed. Endogenous concentration range of TPX2 (30-100 nM) indicated. **(C)** Schematic of method used to determine soluble concentration of GFP-TPX2 at various total concentrations in the presence of BSA and tubulin. Total concentration (squares) is indicated on the Y-axis and the corresponding soluble pool measurement (diamonds) is indicated on the same plane. Mean soluble pool concentrations of three replicate experiments shown. Phase boundary value is the mean SEM of all soluble pool replicates. **(D)** Oblique TIRF images of only Cy5-labeled tubulin condensed with GFP-TPX2 (not shown) either in solution (top panel) or on a pre-formed MT (lower panel – MT not shown) at concentrations shown. Images displayed at matched brightness and contrast. Enhanced contrast of 10nM and 25nM images shown in shown in box. Scale bar, 2µm. **(E)** Quantification of relative tubulin signal from TPX2-tubulin co-condensates in solution (blue curve) and on microtubules (purple curve) at concentrations shown in (E). Mean and SEM of three replicate experiments (error bars) shown. Endogenous concentration range of TPX2 (30-100 nM) indicated. **(F)** Oblique TIRF images (larger fields of view than shown in Fig. 2A) of pre-formed MTs (stabilized with GMPCPP and labeled with Alexa568) in the presence of GFP-TPX2 and Cy5-labeled tubulin (upper panel), GFP and Cy5-labeled tubulin (middle panel), or GFP-TPX2 and Cy5-lableled BSA (lower panel). Note that tubulin (Cy5-labeled) does not bind to MTs (Alexa568-labeled) unless GFP-TPX2 is present. Contrast is maximized in these images. All proteins at equimolar concentration (100 nM). Scale bar, 3µm.

**Fig. S4.**
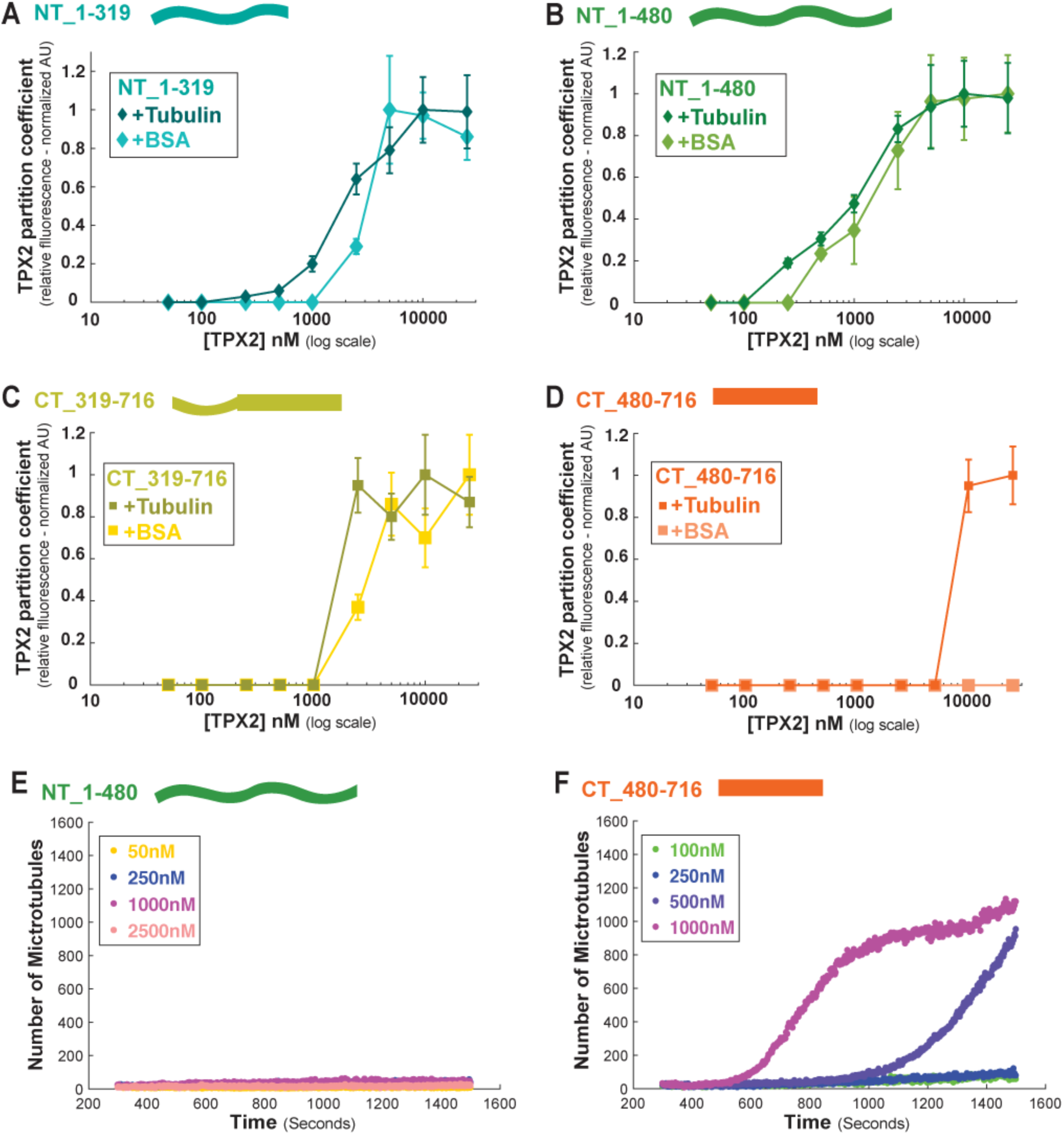
Partition coefficients of TPX2 truncation constructs and their MT nucleation ability. Partition coefficients of GFP_TPX2 in +BSA and +tubulin conditions for the constructs **(A)** NT_1-319, **(B)** NT_1-480, **(C)** CT_319-716, **(D)** CT_480-716. Mean values with ±1 SD as error bars shown. At least 100 condensates per concentration and condition were analyzed. Total number of MTs generated over time for **(E)** NT_1-480 TPX2 and **(F)** CT_480-716 TPX2. Measurements taken at various concentrations of TPX2 (shown in figure).

**Fig. S5.**
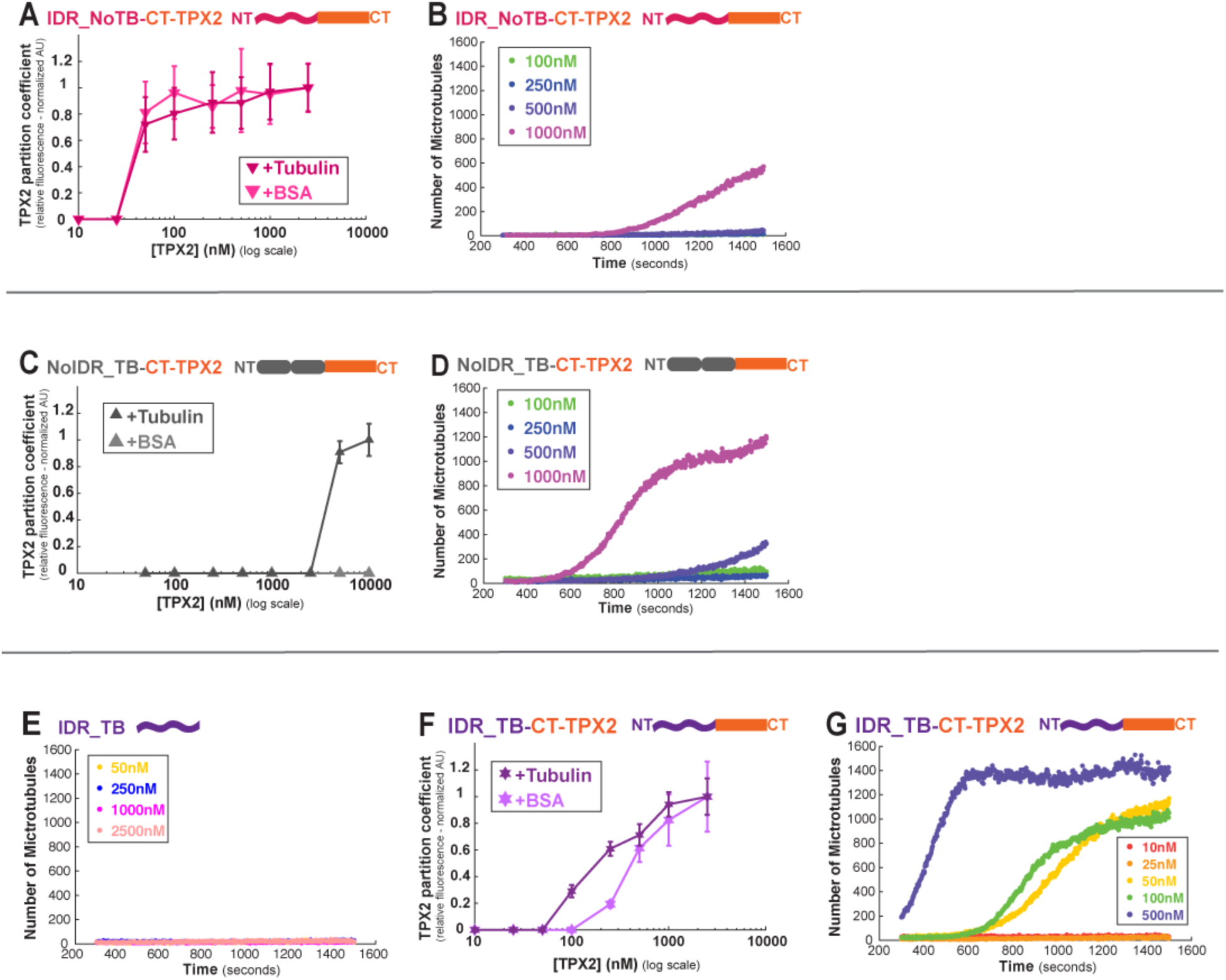
Partition coefficients and MT nucleation rate curves of TPX2 chimera constructs. **(A)** Partition coefficients of GFP_TPX2 in +BSA and +tubulin and **(B)** total number of MTs generated over time for IDR_NoTB-CT-TPX2. **(C)** Partition coefficients of GFP_TPX2 in +BSA and +tubulin and **(D)** total number of MTs generated over time for NoIDR_TB-CT-TPX2. IDR_NoTB-CT-TPX2. **(E)** Partition coefficients of GFP_TPX2 in +BSA and +tubulin and **(F)** total number of MTs generated over time for IDR_TB-CT-TPX2. **(G)** Total number of MTs generated over time for IDR_TB. For partition coefficient graphs mean values with ±1 SD as error bars shown and at least 100 condensates per concentration were analyzed. For both types of graph, measurements were taken at various concentrations of TPX2 and are shown in figure.

**Fig. S6.**
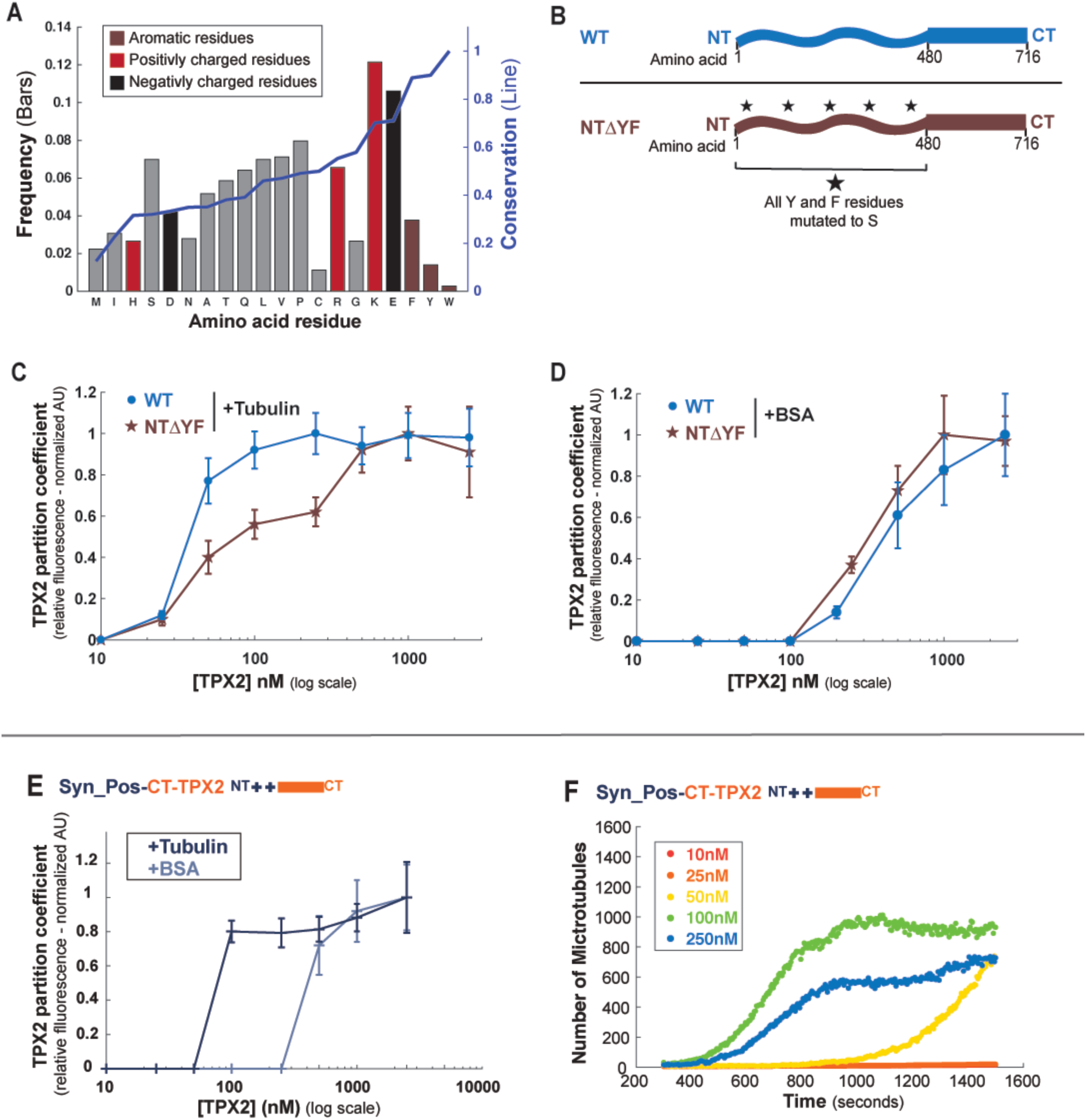
Positively charged residues in TPX2 drive co-condensation with tubulin. **(A)** Graph of amino acids composition of *Xenopus* TPX2. Bars (black borders - left y-axis) displays relative amino acid frequency and line (blue - right y-axis) shows percent conservation of each amino acid type relative to Human TPX2. Amino acids are ordered from least to most conserved and colored (see key). Note that aromatic residues are the most conserved amino acid type, followed by electrostatic residues that are highly conserved and abundant (Lysine –K – being the most abundant). (**B)** Schematic of full length and NTΔYF constructs. Partition coefficients of GFP TPX2 for full length and NTΔYF shown (**C)** +Tubulin and (**D)** +BSA. **(E)** Partition coefficients of GFP_TPX2 in +BSA and +tubulin and **(F)** total number of MTs generated over time for Syn_Pos-CT-TPX2. For partition coefficient graphs mean values with ±1 SD as error bars shown and at least 100 condensates per concentration were analyzed. For both types of graph, measurements were taken at various concentrations of TPX2 and are shown in figure.

**Movie S1.**

GFP_TPX2 condensates falling and fusing on coverslips movie corresponds to Fig. S1

**Movie S2.**

Mono-dispersed GFP_TPX2 (green) localizing to emerging MT fan networks that are marked with Cy5-tubulin (red) and the plus-tip tracking protein EB1 (Blue). Reaction in *Xenopus* egg cytosol treated. Frames were acquired every 10 seconds. Movie corresponds to Fig. 1H.

**Movie S3.**

GFP_TPX2 and Cy5-tubulin co-condensates in *Xenopus* egg cytosol treated with nocodazole to prevent microtubule polymerization. Frames were acquired at the fastest possible rate (one frame per 0.1 seconds) but the rapid dynamics of the co-condensates often lead to an offset in the overlap of their signal (merge channel). Movie corresponds to Fig. 1I.

**Movie S4.**

TPX2-mediated branching MT nucleation in *Xenopus* meiotic cytosol at indicated concentrations of TPX2. Cy5-labeled tubulin (red) and EB1-mCherry (green) highlight microtubules and growing microtubule plus ends, respectively. Movie corresponds to Fig. 2C-E.

**Movie S5.**

TPX2-mediated branching MT nucleation in *Xenopus* meiotic cytosol at indicated concentrations of NT_1-480 TPX2 (top row) and CT_480-716 TPX2 (bottom row). Cy5-labeled tubulin (red) and EB1-mCherry (green) highlight microtubules and growing microtubule plus ends, respectively. Movie corresponds to Fig. 3D and S4E-H.

**Movie S6.**

TPX2-mediated branching MT nucleation in *Xenopus* meiotic cytosol at indicated concentrations of IDR_NoTB-CT-TPX2 (top row) and NoIDR_TB-CT-TPX2 (bottom row). Cy5-labeled tubulin (red) and EB1-mCherry (green) highlight microtubules and growing microtubule plus ends, respectively. Movie corresponds to Fig. 4C and S5B and D.

**Movie S7.**

TPX2-mediated branching MT nucleation in *Xenopus* meiotic cytosol at indicated concentrations of IDR_TB-CT-TPX2 (top row) and Syn_Pos-CT-TPX2 (bottom row). Cy5-labeled tubulin (red) and EB1-mCherry (green) highlight microtubules and growing microtubule plus ends, respectively. Movie corresponds to Fig. 4C and S5F and S6F.

**Movie S8.**

TPX2-mediated branching MT nucleation in *Xenopus* meiotic cytosol at indicated fold excess of importins-α/β. Full-length TPX2 at final concentration of 100nM. Cy5-labeled Tubulin (red) and EB1-mCherry (green) highlight microtubules and growing microtubule plus ends, respectively. Movie corresponds to Fig. 5B-C and E.

